# Vector Competence of Temperate and Tropical Mosquito Species for Babanki virus

**DOI:** 10.1101/2025.04.14.648682

**Authors:** Mathilde Ban, Virginie Geolier, Elisabeth Ferquel, Martin Faye, Moussa Moise Diagne, El Hadji Ndiaye, Gamou Fall, Mawlouth Diallo, Dimitri Lavillette, Valérie Choumet

## Abstract

Babanki virus (BBKV), an alphavirus belonging to the Western Equine encephalitis complex, was isolated from *Mansonia africana* mosquitoes in 1969 but is not characterized yet. Transmitted to humans and animals through the bite of an infectious female arthropod, it is associated with febrile illness, rash, and arthritis similarly to Sindbis virus to which it is related. No specific treatment or vaccine is available. To investigate any risk of propagation, our study investigated the vector competence of several mosquito species towards BBKV. For this purpose, laboratory mosquito colonies from metropolitan France: *Culex pipiens* (Paris), *Aedes albopictus* (Nice) as well as tropical area: *Aedes albopictus* (Saint-Benoît, La Réunion) and *Culex quinquefasciatus* (Slab strain), *Aedes aegypti* (PAEA) were exposed to an artificial blood meal infected with BBKV molecular clone. Midguts, legs, salivary glands and saliva were collected from individual mosquitoes of different strains or species at various time points to assess the infection status, viral replication, and infectiousness. At 7 days post viral exposure, infection rates exceeded 70% in *Aedes* and 40% in *Culex* species. They remain above 50% except for *Culex quinquefasciatus* which decreases to 35% at 14 days post viral exposure. Using genome and protein detection, the dissemination rates are above 50% except for *Aedes albopictus* (Nice) at 7 days post viral exposure. Infection assays using VeroE6 cells indicated that BBKV replicative viral particles were found in the saliva of the different populations tested. Our data demonstrate that different mosquitoes’ species from both temperate and tropical areas are competent to transmit BBKV suggesting an emergence risk that could trigger an outbreak in humans.

**Author Summary:** Babanki virus (BBKV) is a non-characterized arbovirus identified in 1969 in Cameroon. BBKV is closely related to Sindbis virus which causes symptoms like febrile illness, rash, and arthritis in humans and animals. Both are alphaviruses within the Western Equine Encephalitis complex. BBKV is identified only in the African continent so far. Despite its potential to cause disease, no effective treatments are available. This study investigates the vector competence of various mosquito species from temperate and tropical regions, to transmit BBKV. After mosquitoes were fed an infectious blood meal, various organs were collected at different time points to assess the infection status, viral replication, and infectiousness. The results showed that BBKV infection occurred in all tested mosquito species and persisted over time, with viral particles detected in mosquito or that can potentially be transmitted to vertebrate hosts via mosquito saliva. These findings highlight the broad vector competence of mosquitoes from diverse climatic regions for BBKV, indicating its potential risk for emergence and outbreaks in different regions of the world. The study emphasizes the need for enhanced surveillance and preventive measures to mitigate the potential threat posed by this emerging virus.

## Introduction

Alphaviruses have a worldwide distribution, inhabiting all continents except Antarctica (1).They are geographically restricted on preferred ecological conditions, reservoir host and vector species but continue to move around the globe and colonize new areas (2,3). Alphaviruses are maintained in transmission cycles involving both hosts and vectors. Vectors include mosquitoes, ticks, lice and cliff swallow bugs (4,5). The host-range varies from humans and non-human primates, to horses, birds, amphibians, reptiles, rodents, pigs, sea mammals and salmonids. They are categorized into either the Old World or New World group based upon the area in which they were initially found and the disease they cause (6).New World alphaviruses are associated with encephalitis (Venezuelan, Western and Eastern equine encephalitis virus). On the other hand, Old World *Alphaviruses* are responsible for arthritogenic disease (Semliki Forest virus, Sindbis virus, Ross river virus (RRV) or Chikungunya virus (CHIKV)). Few exceptions exist with Sindbis virus (SINV) that is geographically found in the Old World but with a genome that is closer to the Western equine encephalitis (WEE) complex. SINV symptoms can be both encephalitic and arthritogenic (7,8). SINV is geographically widespread and has been found in Africa, Europe, Asia, and Oceania, also designated as Babanki, Ockelbo, Karelian fever, Kyzylagach, or Sindbis-like virus, depending on the location of detection (7,9). The SINV group is divided into six geographically distinct genotypes and antigenically distinct based on their difference in the E2 glycoprotein. The distribution of the genotype SINV-I has been associated with circulation mainly in Africa, Europe and the Middle East. It is this SINV-I genotype that has been associated with human cases (10).

Babanki virus (BBKV) belongs to the Western equine encephalitis complex and is closely related to Sindbis virus (SINV) as previously described (11). The virus was isolated from a pool of 55 adult *Mansonia africana* mosquitoes using a human bait close to high altitude palm trees (*Raphia* species) near the Bambalang and Babanki area in Cameroon (12). The virus was then intracerebrally inoculated to newborn mice and isolated in 1969 (Arbocat, CDC). Subsequent isolations were made in *Anopheles, Aedes, Culex*, *Eretmapodites, Coquillettidia,* and *Mansonia* species throughout much of sub-Saharan Africa and Madagascar (10,13–17). BBKV has been isolated from humans in Senegal, Cameroon, Madagascar and the Central African Republic (6,18) (CRORA, 2000). Recently, neutralizing antibodies were also found in Ugandan bats confirming its circulation (19). BBKV has been associated with human febrile illness accompanied by rash and arthritis. The virus is already spread by mosquitoes and ticks (ArboCat, CDC) from the west to the east coasts of Africa through Central Africa, which highlights its potential to expand and cause outbreaks in new geographic areas.

Like other members of the Togarividae family, BBKV possesses an unsegmented, positive-sense, single-stranded RNA genome approximately 11.7 kilobases in length. This genome encodes both structural and non-structural proteins. The virus has a spherical nucleocapsid about 40 nm in diameter with icosahedral symmetry (T=4), enveloped by a lipid membrane adorned with glycoprotein spikes, resulting in virion particles roughly 70 nm in diameter [https://web.archive.org/web/20070703230421/http://phene.cpmc.columbia.edu/ICTVdB/73001B24.htm]. The strain Y-251 isolated from *Mansonia africana* mosquito species in 1969 in Cameroon had at least 7 different amino acids compared to the HR strain of SINV prototype AR 339 (ratio between synonymous and nonsynonymous substitutions of 3.7). Y-251 is part of a subcluster of Sindbis virus – genotype I that contains isolates from Africa (AR18132, AR18141 from South Africa, MP684 from Uganda and Y-251 from Cameroon) (9). Comparison of the structural and the non-structural polyprotein sequences of SINV MP762-UG-2019 to sequences deposited in Genbank using Basic Local Alignment Search Tool (BLAST) resulted in maximum nucleotide identities of 99.2 and 99.1% to a SINV isolate named Babanki virus (HM147984), respectively (15). This cluster is different from the one in Europe or the Middle East. In 2011, Forrester *et al.*, indicated that the sequence of BBKV is a subtype of SINV as part of the WEE antigenic complex using the full genome alignment of both ORFs. So BBKV was classified as a “new world” virus.

In this study, we aim to evaluate the vector competence of several mosquito species from the *Aedes* and *Culex* genera, representing both tropical and temperate regions. Both genera are recognized vectors of alphaviruses (20–22). *Aedes aegypti*, predominantly found in tropical areas, will be represented by a laboratory-maintained colony. *Aedes albopictus*, although originally native to tropical regions, has successfully adapted to temperate climates. To reflect this geographical variation, we will use colonies originating from La Réunion Island (Indian Ocean) and from southern France, representing tropical and temperate environments, respectively. For *Culex* species, colonies from Paris and southern metropolitan France will be used to capture the diversity of temperate populations.

Although Sindbis virus (SINV) is known to cause large outbreaks, the emergence potential of the Babanki virus (BBKV) subtype remains largely unknown (18). While BBKV shares similarities with other alphaviruses such as SINV, its unique biological characteristics, transmission dynamics, and public health impact have yet to be elucidated. It is therefore essential to conduct research that deepens our understanding of BBKV and provides tools to assess its potential for emergence in new geographic regions.

The objective of this study was to characterize the emergence potential of the African alphavirus Babanki by evaluating the vector competence of multiple mosquito genera and species from diverse regions. To this end, we constructed a molecular clone of the virus based on the sequence of the ARY168 strain. Our findings demonstrate, for the first time, that both temperate and tropical mosquito species are competent vectors for BBKV, highlighting its potential for emergence across a range of ecological settings. These results underscore the critical need for ongoing surveillance and vector control measures to mitigate the risk of future outbreaks.

## Methods

### Cells

VeroE6 cells (African green monkey epithelial cells) were maintained in culture with Dulbecco’s Modified Eagle Medium (DMEM) 1X + GlutaMAX^TM^ (ref. 3166-021, Life Technologies) supplemented with 10% of heat-inactivated fetal bovine serum (FBS) (ref. A15-701, lot A70108-2369, PAA) and 1% of Penicillin-Streptomycin (P/S) (ref. 15140-122, Life Technologies) at 37°C with 5% CO_2_. C6/36 *Aedes albopictus* cells were cultured in Leibovitz’s L15 medium 1X (ref. 11415-049, Life Technologies) with 10% FBS and 1% P/S at 28°c without CO_2_.

### Virus

#### Viral sequence

The full length ARY168 11 724 bp DNA sequence was generated from a virus isolated from *Culex decens* mosquito species (Yaoundé, Cameroon 1970) by Institut Pasteur of Dakar.

### Mapping of ARY168 sequence

Snapgene software was used to find open reading frames (ORF). Non-structural and structural proteins were determined based on the sequence alignment analyses with annotated reference sequences (BBKV HM147984.1 and Sindbis virus NC_001547.1) on Ugene.

### Identification of predicted *E. coli* promoter sequences within BBKV genome ARY168

The Neutral Network Promoter Prediction (NNPP) from the Berkeley *Drosophila* Genome Project (https://www.fruitfly.org/seq_tools/promoter.html) was used to identify predicted *Escherichia coli* (*E. coli*) promotors (ECPs) sequences for cryptic expression of viral protein.

### Add-ons

**A T7 promotor** sequence was added at the 5’ UTR. A termination sequence including a hepatitis delta virus (HDV) antigenomic ribozyme and a simian virus (SV 40) polyadenylation signal (pA) (SV40pA) was placed at the 3’ UTR.

### Fragmentation

Mutated added-on BBKV sequence was divided into 4 fragments with a pace of approximately 3.000 nt framed by unique restriction sites.

### Plasmid

High copy number plasmid with ampicillin resistance gene pUC18 was chose for cloning. The initial multiple cloning site (MCS) of pUC18 was changed to ClaI-SpeI-PsiI-XhoI and NotI (32 nt) to a final pUC18_MSC_BBKV plasmid. To this aim, small oligos were hybridized at 95°C for 5 min and then cooled to room temperature. After purification, hybridized DNA was ligated to the open pUC18 plasmid with T4 ligase (ref. M0202, NEB).

### Cloning

The 4 synthetics genes were cloned into the pUC18_MSC_BBKV using both T4 ligase (ref. M020, NEB) or the recombinases In-Fusion® HD Cloning Kit (ref. 638910, Takara)/Ezmax One-Step Cloning Kit (ref. 24303-2, Tolobio) strategies. Cloned plasmids were transformed into Top10 or JM110 competent bacteria. Mini cultures with DNA extraction were performed prior to maxi cultures and DNA extraction. Molecular clone was double checked by digestion restriction and by sequencing with BioSun company (China).

### *In vitro* transcription

Full length BBKV molecular clone DNA was linearized with NotI. After extraction with phenol-chloroform, Linear DNA was precipitated with ethanol and dissolved in RNAse free water. The *in vitro* transcription was achieved using mMESSAGE mMACHINE T7 (ref. AM1344, Invitrogen) kit following the manufacturer’s instructions. The transcript mRNA was precipitated by LiCl solution. RNA transcript was checked by electrophoresis and quantified.

### Electroporation

10 μg of full-length *in vitro* transcribed BBKV mRNA was electroporated into VeroE6 cells using the GenePulser XCell electroporation system (Bio-Rad) with 4 mm cuvettes and the following setting: 270V, 975 μF, ∞ Ω, 4 mm, 1 pulse. Cells were incubated at 37°C with 5% CO2 in a BSL-3 facility. After 4 hours post-electroporation, the cell’s supernatant was removed and replaced with 2% FBS fresh cells media. The culture supernatant was collected 30h post electroporation when cytopathic effect (CPE) was observed, clarified by centrifugation, and re-amplified by infection of fresh VeroE6 or C6/36 cells. Cultured supernatant was harvested when CPE were present around 30 hrs post amplification), clarified by centrifugation, aliquoted and stored at −80°C.

### Titration by plaque assay

The produced virus was 10-fold diluted until 10^-10^. VeroE6 cells at 90% confluence were infected with 200 μL of virus dilutions in DMEM supplemented with 2% of FBS in the BSL-3 environment. After 1 hour, the inoculum was removed and replaced with 2% final concentration carboxymethyl cellulose (CMC) with DMEM and 2% FBS 1% P/S media. After 24 hours at 37°C with 5% CO2, cells were washed from the CMC polymer mix with PBS and fixed with a 4 % PFA solution during 30 min. After washing the PFA with PBS, the plate was stained with crystal violet 0.1% solution. Plaque per dilution were counted to compute the viral titer that is expressed as plaque forming unit per ml (pfu/ml).

### Antibodies

Noncommercial mouse polyclonal antibodies against Sindbis virus were used as primary antibodies. Mouse polyclonal antibodies against *Aedes aegypti*, *Aedes albopictus* and *Culex pipiens* salivary glands proteins were used as primary antibodies.

Calnexin polyclonal antibody purified from rabbit serum (ADI-SPA-865, Enzo Life Sciences) was used as a primary antibody to detect housekeeping genes in mammals.

Both Goat anti-Mouse IgG (H+L) HRP conjugated (Cat#172-6516 Bio-Rad) and Goat anti-Rabbit (Cat#172-6515 Bio-Rad) were used as a secondary antibody for chemiluminescence. For fluorescent microscopy, Alexa Fluor 488 AffiniPure Goat anti-mousse IgG (H+L) Jackson ImmunoResearch (ref. 115-545-003) was used.

### Mosquitoes strain

*Aedes aegypti* (PAEA) strain is a laboratory colony (more than 400 lab-generations) isolated in 1960 in French Polynesia. *Aedes albopictus* Nice strain was collected in Nice, South of France in 2011. *Aedes albopictus* Saint-Benoit strain was collected in Saint-Benoit, La Réunion Island in the Pacific Ocean in 2012. *Culex pipiens pipiens* strain was collected in Paris in 2017. *Culex quinquefasciatus* (Slab strain) was kindly donated by Mylène Weill from the Institute of Evolution Sciences of Montpellier (ISEM) (University of Montpellier, CNRS, IRD, EPHE, CIRAD, INRAP).

### Mosquito breeding

From the egg to the female adult for blood meal, it takes around 4 weeks. Every stage of the mosquito development takes place in a room at 27°C, 70% of relative humidity with a 12 hour photoperiodicity. *Aedes* eggs are stored on dry blotting forest green paper up to 3 months whereas *Culex* eggs cannot be stored. Mosquito eggs are put in 2 L of non-limestone water with cats and fish dry food. Depending on the species, dry brewer’s yeast can be added to boost the development. Larvae from stage 1 to 3 are reared in the same condition in netted-covered flat bucket. Pupae are sorted and placed in a netted cage before emergence. Adult male emergence happens few days before the female one. Adult mosquitoes are fed *ad libitum* on 10% sucrose solution. Two blood meals at 3 days apart are necessary for the females to lay eggs on water (*Culex*) or blotting paper (*Aedes*). For this purpose, mice are anesthetized with an injectable solution of ketamine/xylazine mix and ventral sided placed on the top of the netted cage for 20 min. The blood meal feeding on living animals is approved by the ethics committee of Institut Pasteur C2EA n°89 under the agreement 2014-0068.

### Exposure to infectious blood meals

Week-old female mosquitoes were sorted and starved 24 hours prior to the infectious blood meal. On the infection’s day, fresh rabbit blood was collected on pre-heparinized 15 mL tubes and washed 3 times with PBS. The infectious blood meal was constituted in the BSL-3 facility and consists in one third of viral suspension (final concentration of 1.10^8^ PFU/mL), two thirds of rabbit erythrocytes and adenosine triphosphate (ATP) as a phagostimulant to a final concentration of 10 mM. *Aedes* species were allowed to feed for 20 minutes through a collagen membrane (*Aedes aegypti*) or swine intestine (*Aedes albopictus*) covering electric feeders maintained at 37°C (Hemotek system). *Culex* species were allowed to feed for 30 minutes directly on blood-soaked cotton at 37°C in an incubator with 5% CO_2_ and without light. After feeding, mosquitoes are anesthetized at 4°C. Blood-fed female mosquitoes are selected on ice and transferred into netted-cardboard boxes with *ad libitum 10%* sucrose solution and kept in an incubator at 28°C (*Culex quiquefasciatus* and *Aedes aegypti*) and 24°C for the other temperate species with 80% of humidity and a 12 hour photoperiodicity incubator for a maximum duration of 14 days.

### Samples collections/Mosquito dissection

**In the BSL-3 facility,** mosquitoes were anesthetized on ice at 7 and 14 days after the infectious blood meal exposition. They went through a 70% ethanol bath and a PBS one. They were dissected in a drop of PBS under a magnifying glass with micro dissecting needles and micro dissecting needle holders (Fine Science Tools, Foster City, USA) and tweezers (#11252-23, Dumont tweezer - Style 5 - Dumoxel). For RNA extraction, midguts, legs and salivary glands were collected from the same mosquito and separately homogenized in 350 μL of RA1 lysis buffer from Nucleospin RNA kit (ref. 740955.50 Macherey-Nagel) with 1% of *β*-Mercaptoethanol (M6250, Sigma-Aldrich) and sterile glass beads (ref. 152018, Dutscher) and then stored at −80°C. For infectious particles quantification, midguts and salivary glands were collected from the same mosquito and separately homogenized in 500 μL of DMEM 2% SVF 1% P/S and 1X ATB/ATF. Homogenization was performed by the Precellys system (Bertin Technologies) at 10 000 *x g* 15 sec twice with a 10 sec break and kept at −80°C. For viral proteins detection, midguts and salivary glands from the same mosquito were collected and individually stored in 10 μL of RIPA 2X buffer (ref. RB4476, BioBasic Canada) at −20°C.

For the saliva samples, mosquito legs were first removed. Then, the mosquito proboscis was inserted in 10 μL tips pre-filled with PBS. From this saliva collection per mosquito, the sample is split in three. Saliva samples for RNA extraction were lysed in 100 μL of RA1 lysis buffer and 2 μL of reducing agent TCEP (Tris(2-carboxyethyl) phosphine) from Nucleospin RNA XS kit (ref. 740902.50 Macherey-Nagel). Samples for viral protein detection were mixed with a 1:1 ratio of RIPA 2X buffer (ref. RB4476, BioBasic Canada) and kept at −20°C. The remaining volume was used with DMEM 2% SVF 1% P/S and 1X ATB/ATF for quantification of infectious particles.

### Viral RNA quantification/RT-qPCR analysis

*RNA extraction.* After lysis, total RNA was extracted using the Nucleospin RNA kit (ref. 740955.50 – Macherey-Nagel) following manufacturer’s instructions. *Reverse transcriptase (RT) and quantitative PCR (qPCR).* Viral RNA quantification was performed using 2 μL of extract RNA with the *Power* SYBR^TM^ Green RNA-to-CTTM *1- Step* Kit (ref. 4389986, ThermoFisher Scientific) following the manufacturer’s instructions. Specific primers were used to amplify the viral RNA:

nsp2-Forward: AAGCGATGCGTTAAGAAGGA
nsp2-Reverse: ACTTGATGATAGCTGACTTGCC.

The QuantStudio 6 Flex Real-Time PCR System (Applied Biosystem) monitors the fluorescent signal produced by the SYBR^TM^ Green which binds double-stranded DNA. The thermal cycling condition are: reverse transcription for 30 min at 48°C; 10 min at 95°C and then 40 cycles of 15 sec at 95°C followed by 1 min at 60°C. Quantity of RNA copies per mL of sample was extrapolated from the standard curve generated from the 10-fold serial dilution of the *in vitro* transcript mRNA previously quantified by the ND-1000 Spectrophotometer (Thermos Fisher Scientific).

### Viral protein detection/Blot analysis

#### Western Blot

After thawing, midguts and salivary glands samples collected in RIPA buffer were centrifuged at maximum speed during 5min at 4°C. Then samples were denatured in 1X NuPage LDS Buffer (ref. NP0008, ThermoFisher Scientific) during 5 min at 95°C and loaded to a precast Nu-Page 4-12% Bis-Tris Gel (ref. NP0322BOX, ThermoFisher Scientific). Proteins were separated by electrophoresis at 60 mA and transferred thanks to a Trans-Blot^®^ TurboTM Mini PVDF Transfer Packs (ref. 1704156, Bio-Rad) with the Trans-Blot^®^ TurboTM System (Bio-Rad).

#### Dot blot

An Immuno-blot PVDF membrane (ref. 1620177; Bio-Rad) was activated by submerging it in 100% ethanol and rinse it with PBS. Then, we applied 2 μL of each saliva sample in the RIPA buffer in one square. The blot was then air-dried for 5 to 10 min. After blocking with PBS - Tween^®^20 0.1% (ref. P7949, Sigma Aldrich) - Skim Milk 5% (ref. 232100, BD) during 1 hour at room temperature under agitation, the PVDF membrane was incubated overnight at 4°C with primary antibody diluted in blocking buffer under agitation. Finally, the membrane was incubated with the secondary antibody diluted in the blocking buffer during 1 hour at RT under agitation. Proteins were revealed using Pierce^TM^ ECL Western Blotting Substrate (ref. 32106, ThermoFisher Scientific) and MyECL^TM^ Imager (ThermoFisher Scientific).

### Results interpretation

To determine vector competence of each mosquito species, infection, dissemination and transmission rates are computed according to the titration by plaque assay results (from the midguts and salivary glands in DMEM buffer) and the following formulas:

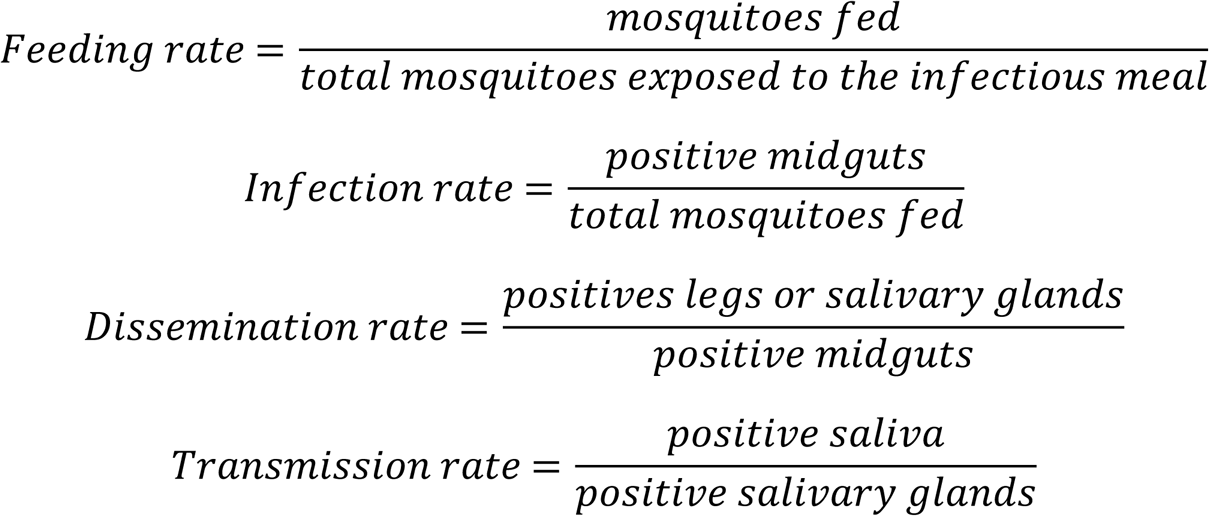

### Biostatistical analysis

Statistical analyses were computed using Prism 6 and SimStat software. Nonparametric Kruskal-Wallis and Mann-Whitney tests were used to test differences between groups (correction of Tuckey). P-value inferior at 0.05 are considered significant following: *: p<0.05; **: p<0.01; ***: p<0.001; ****: p<0.0001.

## Results

### Infectious molecular clone construction

#### Identification of predictive *E. coli* promoter sequences within BBKV genome ARY168

The full-length genome BBKV ARY168 sequence was kindly provided by IP Dakar. Non-structural and structural proteins were determined on the 2 open reading frames (ORF) based on the sequence alignment analyses with annotated reference sequences (BBKV HM147984.1 and Sindbis virus NC_001547.1) on Ugene. Based on the previous work done for Yellow fever and Japanese encephalitis viruses by the team of Pu *et al.*, (23), putative predicted *Escherichia coli* (*E. coli*) promotors (ECPs) sequences for cryptic expression of viral protein were found using the Neutral Network Promoter Prediction (NNPP) from the Berkeley *Drosophila* Genome Project (https://www.fruitfly.org/seq_tools/promoter.html). In total, 14 putative predicted ECPs with a score higher than 0.9 were identified for the BBKV genome ARY168 sequence represented in Table 1.

**Table 1:**
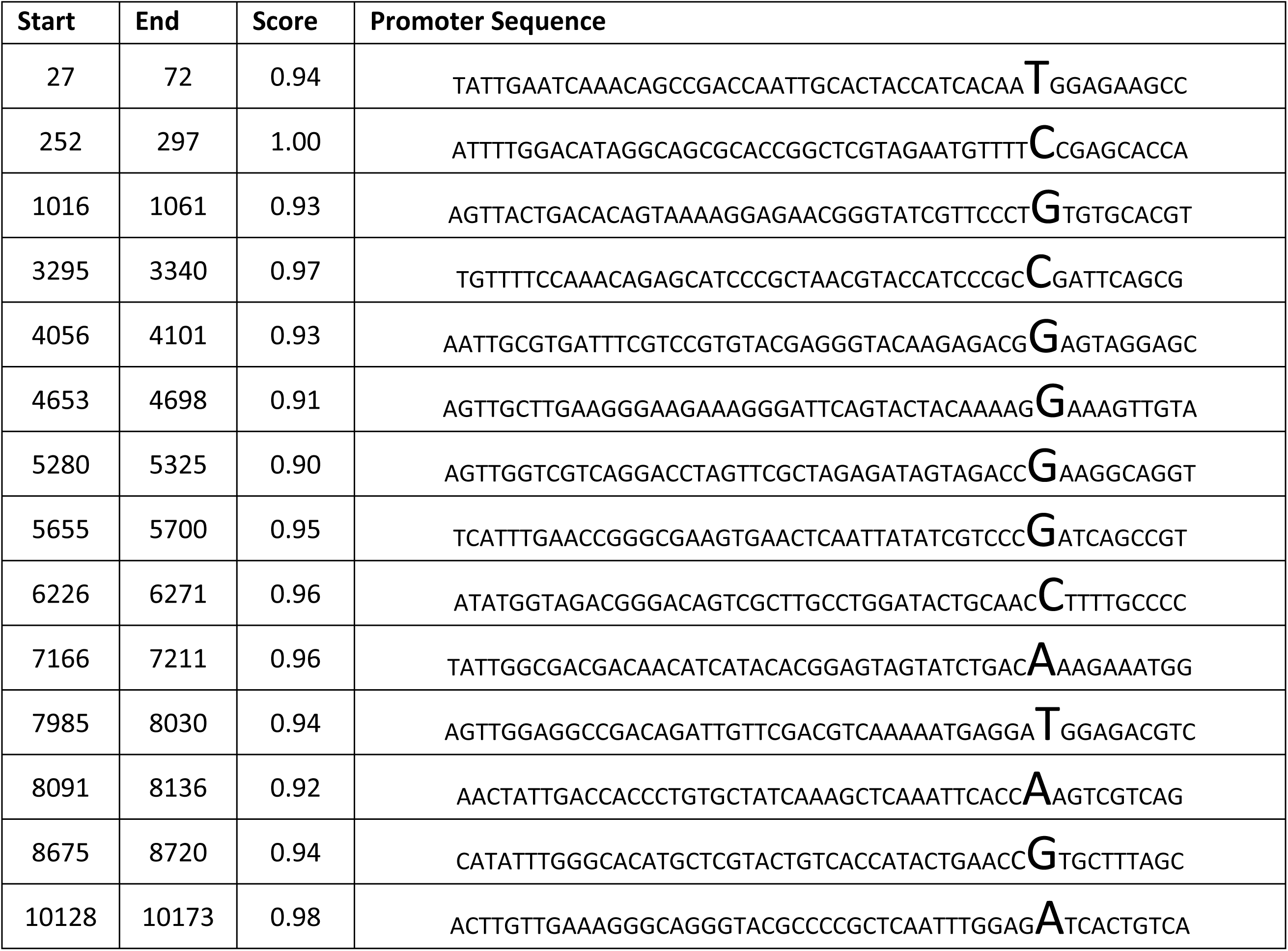
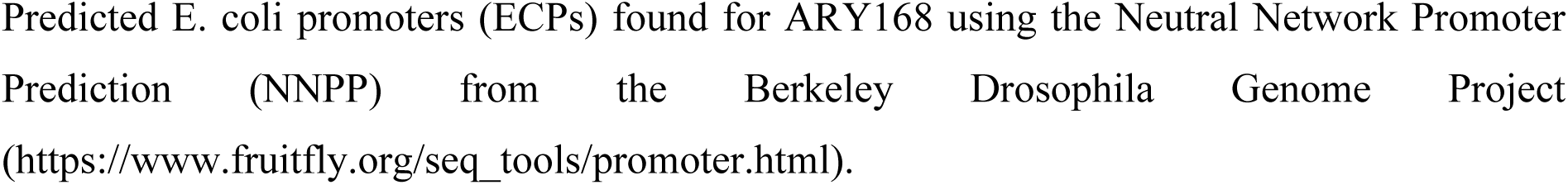
Predicted *E. coli* promoters (ECPs) found for ARY168.

#### BBKV molecular clone construction *in silico*

Silent mutation introduction in the BBKV genome did not affect the amino acid sequence of ARY168 genome. Silent mutations were manually introduced to lower the score to 0, except in the 5’ and 3’ UTR region and the non-coding part between the structural and non-structural protein genes. In total, 6 ECPs scores were lower to non-detectable (ND) keeping the same amino acid sequence (Fig 1.A). These mutations were generated to improve the stability of the BBKV genome in bacteria when amplified.

**Fig 1:**
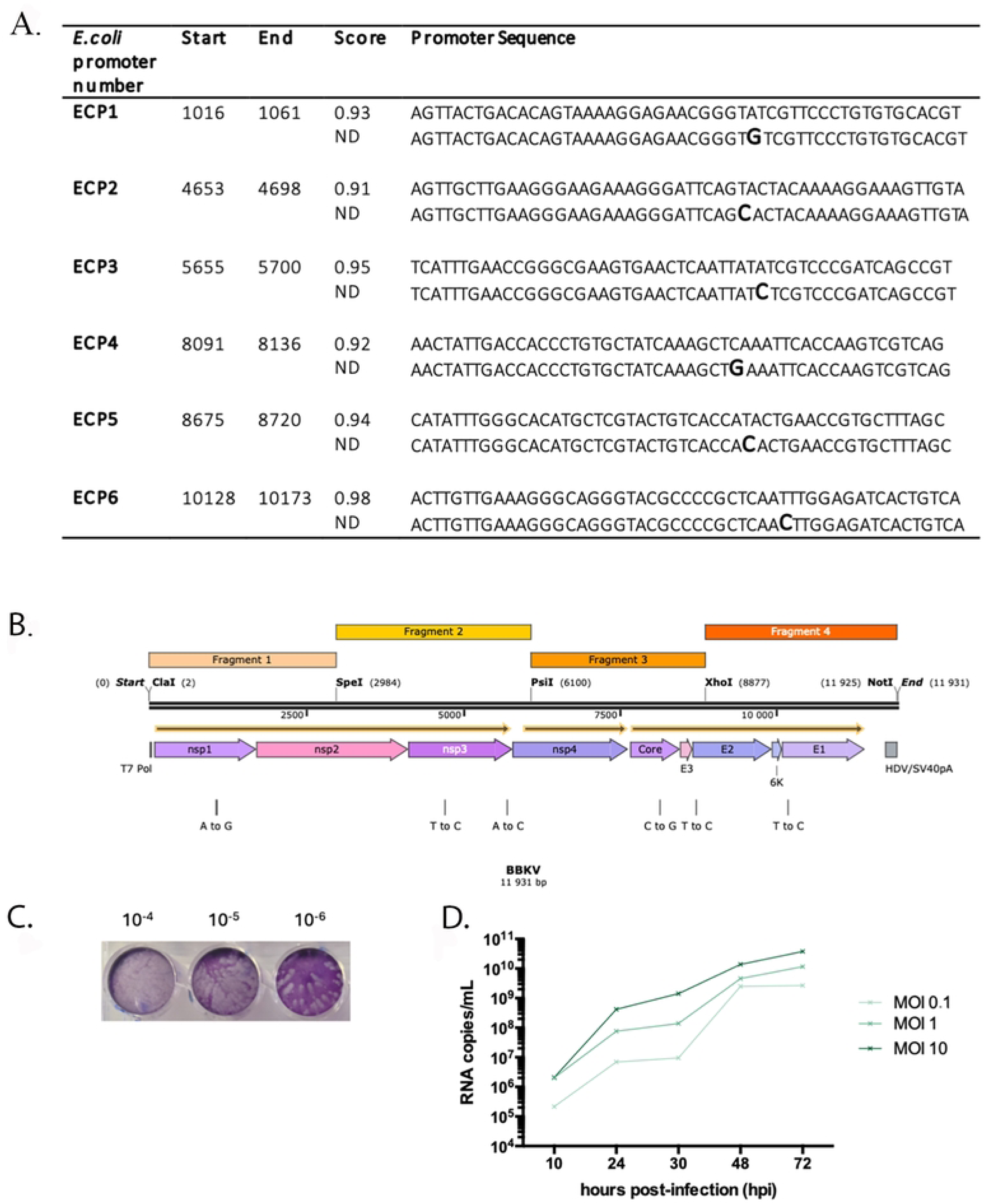
BBKV molecular clone construction and validation. **A.** Silent mutations introduced in ECPs in bold letters. New scores were not detectable (ND) with the NNPP. **B**. Annotated map of mutated BBKV ARY 168 genome with Snapgene and Ugene softwares. C. Observation of Plaque Forming Unit on infected VeroE6 cells at different BBKV dilutions. D. Replication kinetics of BBKV on C6/36 cells at a multiplicity of infection (MOI) of 0.1; 1 and 10. Total RNA was quantified at different time points in cell supernatant.

**Using SnapGene** in silico, a specific T7 promotor sequence was added at the 5’UTR and a termination sequence including a hepatitis delta virus (HDV) antigenomic ribozyme (Perrotta and Been, 1991) and a simian virus (SV 40) polyadenylation signal (pA) (SV40pA) (Hans and Alwine, 2000) was placed at the 3’ UTR. Combination of HDV ribozyme and SV40pA cassette would allow to generate more efficient gene expression (Yamanaka and Xanthopoulos, 2004). **Unique** restriction sites were introduced for the cloning of the mutated full-length BBKV genome that was fragmented into 4 sections of 3.000 nt approximately (Fig 1.B).

#### BBKV molecular clone construction and validation

High copy number pUC18 plasmid contains an ampicillin resistance gene that will be used as a selection gene by ampicillin antibiotic. pUC18 original multiple cloning site (MSC) was replaced by MSC_BBKV (ClaI-SpeI-PsiI-XhoI-NotI) allowing the cloning of BBKV synthetic gene fragments manufactured by Generay Company (China). The final full-length construction was confirmed by digestion restriction and by sequencing analysis. Plaque forming unit from the viral stock on VeroE6 cells (Fig.1.C). Replication kinetics was also tested on C6/36 cells (Fig.1.D).

### Vector competence

#### Mosquito feeding rate

Mosquitoes were exposed to an infectious BBKV blood meal at a viral titer of 1×10^8^ PFU/mL. The viral titer was double checked after the meal exposition by titration assay. The feeding rate was calculated as the proportion of mosquitoes that ingested the blood meal over the number of total mosquitoes exposed to the infectious blood meal (Table 2). Significant variation was observed among mosquito species (p<0), with *Culex quinquefasciatus* exhibiting the highest feeding rate (95%), while *Ae. aegypti* had the lowest (48%). *Ae. albopictus* (Nice and Saint-Benoît) showed similar feeding rates (∼60%), whereas *Culex pipiens* had a moderate feeding rate (82%).

**Table 2:**
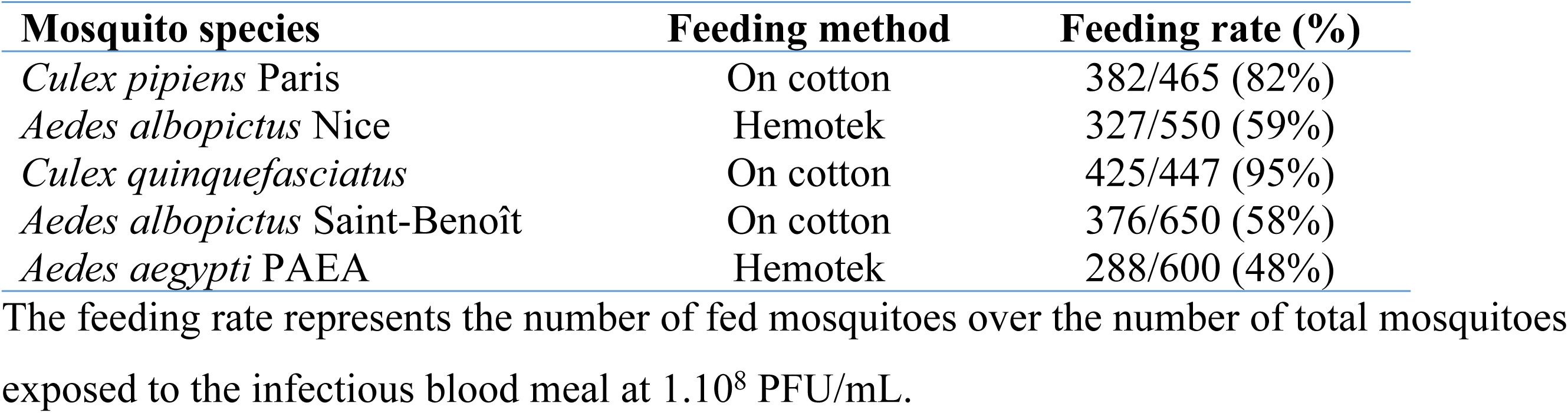
Feeding rate after the exposition to BBKV infectious blood meal.

#### Infection and persistence

At 7-days post viral exposition (dpve), viral RNA was amplified from total RNA extracted from all mosquitoes’ midguts (Fig 2.A). A heterogeneous viral RNA distribution was observed, with *Culex* species having higher copy numbers (>10⁶ RNA copies/midgut) compared to *Aedes albopictus* (Nice) (∼10⁵ RNA copies/midgut). For *Aedes aegypti* (n=30), 3 distinct midgut pools could be identified: one with a high copy number, an intermediate one and a lower one. For *Aedes albopictus* Saint-Benoit, there was only one pool of high copy number of viral RNA copies over 10^7^ RNA copies/ organ.

**Fig 2:**
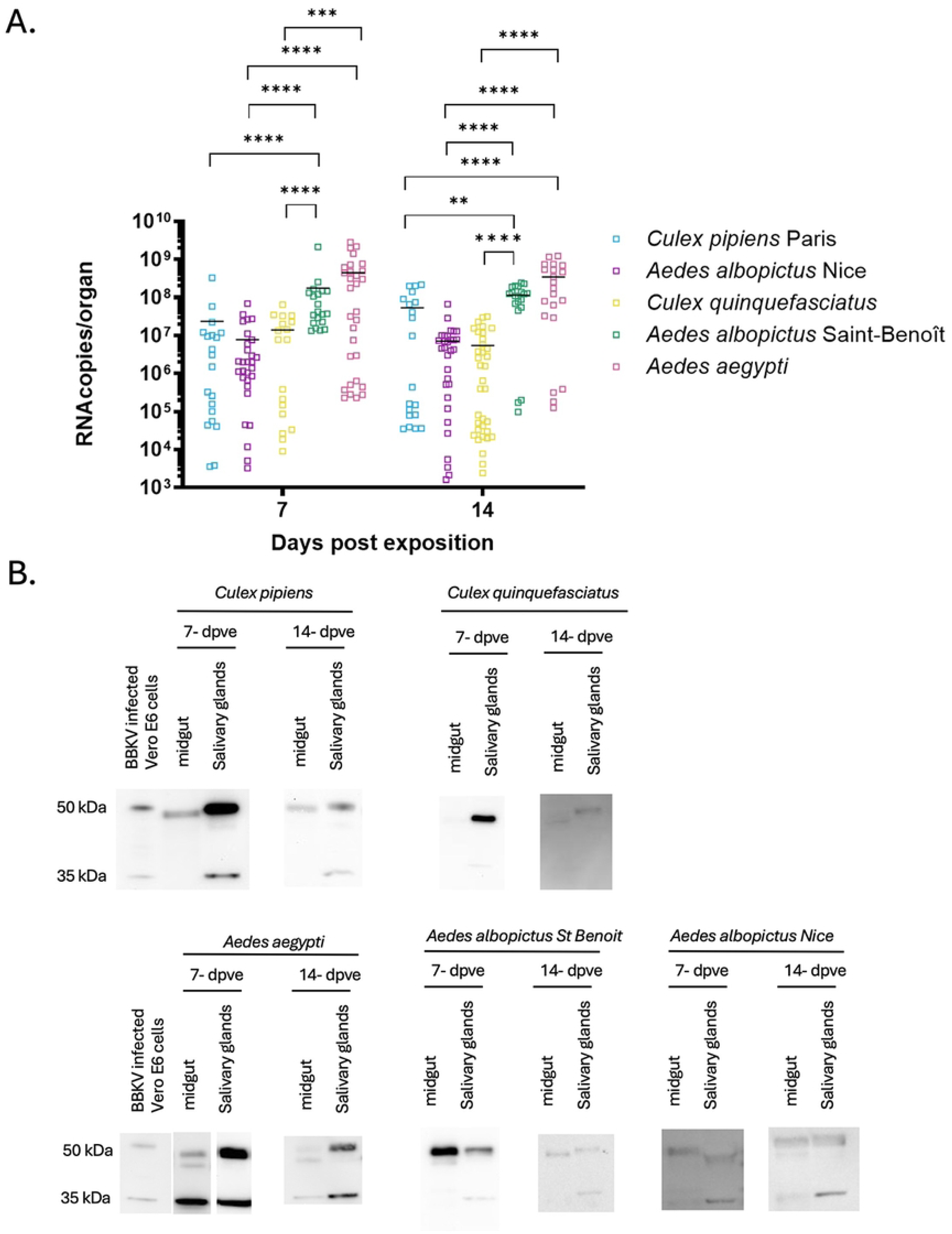
Viral amplification and detection at 7- and 14-dpve. **(A)** Viral RNA amplification for genome quantification by RT-qPCR in mosquito midguts for BBKV. At 7 dpve, n=30 for *Aedes albopictus* Nice and *Aedes aegypti*; n=20 for *Culex quinquefasciatus*, *Aedes albopictus* Saint-Benoît and *Culex pipiens* Paris. At 14-dpve, n= 20 for *Culex pipiens* Paris and *Aedes albopictus* Saint-Benoit; n=22 for *Aedes aegypti*; n=34 for *Aedes albopictus* Nice and n=40 for *Culex quinquefasciatus*. **: p<0.01; ***: p<0.001; ****: p<0.0001 (Mann-Whitney non-parametric test).**(B)** BBKV viral protein detection in the MG and SG. Mosquito organs were collected in the RIPA buffer. After SDS-PAGE electrophoresis and transfer to a PVDF membrane, SINV ascite was used. Bands at 35 kDa represent core protein and 50 kDa preE1-E2.

At 14-dpve, *Culex pipiens* Paris (n=20), *Aedes albopictus* Saint-Benoît (n=20) and *Aedes aegypti* (n=22) had a heterogeneous distribution with a high pool around 10^7^-10^8^ RNA copies/midguts and a lower one at 10^4^-10^5^ RNA copies/midgut. However, *Aedes albopictus* Nice and *Culex quinquefasciatus* midgut viral RNA had a homogeneous distribution ranging from 10^3^ to 10^8^ RNA copies/midguts.

Viral loads at 7- or 14-dpve were non-significantly different within the same species. If we compared the viral loads of the same climate area species, *Culex pipiens* from Paris and *Aedes albopictus* from Nice were not significantly different. On the other hand, tropical species had a significantly different (p<0.0001) viral load in the midgut.

Since viral genome was detectable at 7- and 14-dpve from the mosquitoes exposed to an infectious blood meal, we can suggest that the virus is persistent in the mosquito midgut for the species tested.

Viral structural proteins could be detected in the mosquito midguts at both 7- and 14-dpve (Fig 2.B).

#### BBKV replication in the mosquito midgut and persistence

Infectious particles or replicating viruses were quantified in the midguts of mosquitoes at 7- and 14-dpve (Fig 3). A statistical difference was found between *Aedes albopictus* Nice and *Aedes aegypti* at 7-dpve whereas a high statistical difference was found between viral titers measured in the midguts of the different species at 14-dpve. The viral titer found in *Aedes albopictus* Nice, *Aedes albopictus* Saint-Benoît and *Aedes aegypti* were similar and significantly higher than those of *Culex pipiens* and *Culex quinquefasciatus*. In conclusion, BBKV replicated better in the midguts of *Aedes* species than in those of *Culex* species. We then compared the difference of viral replication between 7-and 14-dpve. A significantly decreasing viral titer was found in *Culex pipiens* and *Culex quinquefasciatus* midguts between the two time points (p<0.04 and p<0.0065 respectively). Interestingly, a significant increase of the viral titer was observed for *Aedes albopictus* Saint-Benoît (p<0.027) whereas similar titers are observed for *Aedes albopictus* Nice and *Aedes aegypti*.

**Fig 3:**
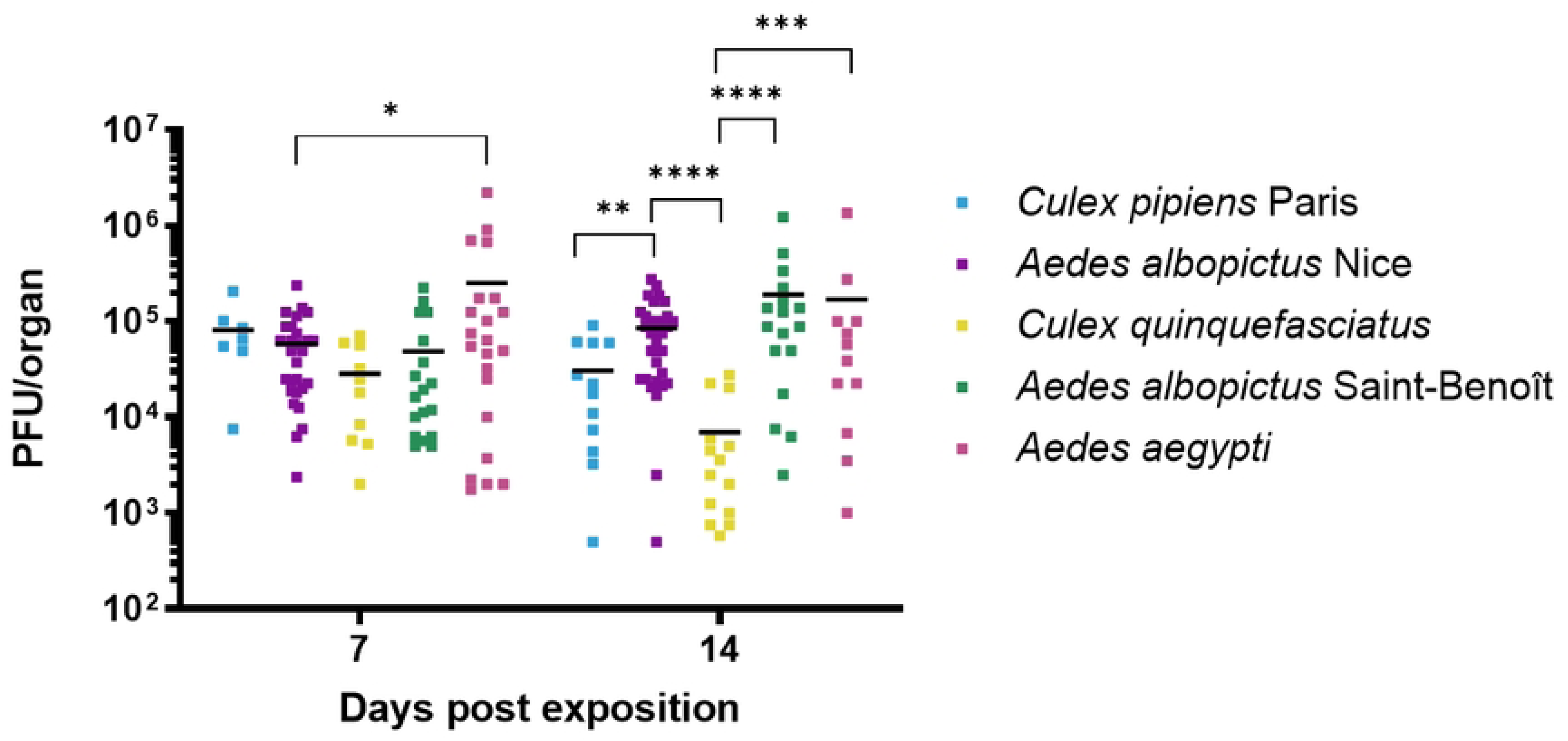
Viral titer in MG at 7- and 14-dpve. After the infected artificial blood meal, MG from the different mosquito species were collected and viral replicative particles were quantified by plaque assay titration on VeroE6 cells. *\*: p<0.05 **: p<0.01; ***: p<0.001; ****: p<0.0001 (Mann-Whitney non-parametric test)*.

The infection rates at 7- and 14-dpve (Table 3) were statistically different according to the different mosquito species (p<0.00003 and p<0.00006 respectively). Regarding the results at 7-dpve, *Culex pipiens* mosquitoes had a lower rate of infection than both populations of *Aedes albopictus* (p<0.0004 for *Aedes albopictus* Nice and for *Aedes albopictus* Saint-Benoît). The rate of infection of the three *Aedes* mosquitoes was not statistically different. The rate of infection of *Culex quinquefasciatus* was statistically like that of *Culex pipiens*. In conclusion, the *Aedes* mosquitoes had a higher infection rate than the *Culex* mosquitoes tested.

**Table 3:**
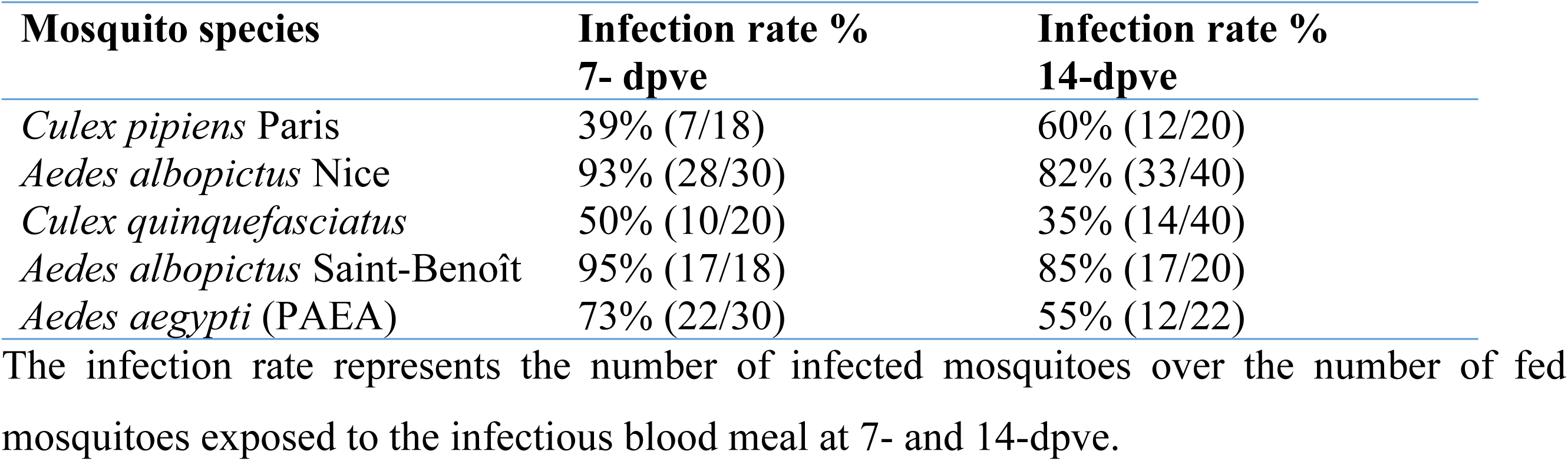
Infection rates at 7- and 14- dpve.

At 14-dpve, the infection rates of *Aedes albopictus* strains, *Aedes aegypti* and *Culex pipiens* were statistically similar. The rate of infection of *Culex quinquefasciatus* mosquitoes dropped and was statistically inferior to that of the two populations of *Aedes albopictus* (p=0.00002 for the European strain and p<0.0003 for the tropical strain).

#### BBKV dissemination to the mosquito leg and persistence

Viral RNA was detected in mosquito legs at 7- and 14-dpve to confirm the BBKV dissemination from the midgut escape barrier and persistence. There was a heterogeneous distribution of the viral RNA in all the species studied: a very high pool and a lower one (Fig 4). As for the midguts, there was no significant difference within each species at 7- and 14-dpve. For *Culex pipiens* from Paris and *Aedes albopictus* from Nice, the viral load in the legs was significantly different (p<0.05) at 14-dpve. There was also a significant difference (p<0.05) between *Aedes albopictus* from La Réunion and *Culex quinquefasciatus* at 14-dpve. This suggests that there was a difference in the way the viral RNA of BBKV persisted in the mosquito legs.

**Fig 4:**
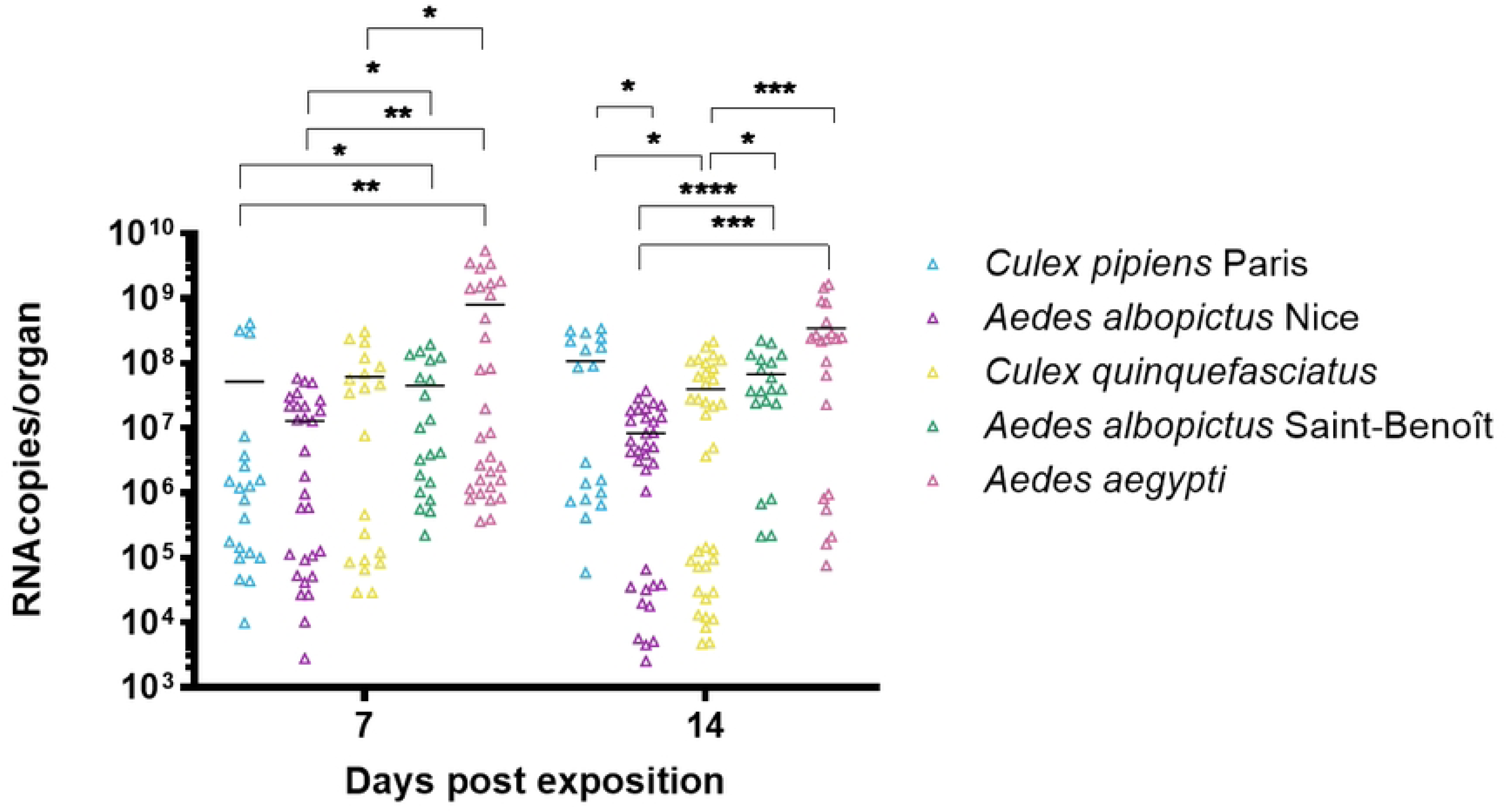
Viral RNA amplification for genome quantification by RT-qPCR in the legs at 7- and 14-dpve. At 7-dpve, n=30 for *Aedes albopictus* Nice and *Aedes aegypti*; n=20 for *Culex quinquefasciatus*, *Aedes albopictus* Saint-Benoît and *Culex pipiens* Paris. At 14-dpve, n= 20 for *Culex pipiens* Paris and *Aedes albopictus* Saint-Benoit; n=22 for *Aedes aegypti*; n=34 for *Aedes albopictus* Nice and n=40 for *Culex quinquefasciatus*. *: p*<0.05; **: p<0.01; ***:p<0.001; ****: p<0.0001 (Mann-Whitney non-parametric test)*

#### BBKV dissemination to the mosquito salivary glands and persistence

Viral RNA was extracted from salivary glands collected on the same mosquitoes for which we previously analyzed midguts and legs at both 7- and 14-dpve and analyzed by RT-qPCR (Fig 5). Viral RNA genome levels were not significantly different in the SG at 7- and 14-dpve within each species. At 7-dpve, viral RNA copy levels in the SG of *Culex pipiens* Paris and *Aedes albopictus* Nice were significantly different (p<0.005). For *Culex quinquefasciatus* and *Aedes albopictus* from La Réunion, the amount of viral RNA copies was not significantly different. At 14-dpve, the amount of RNA copies per pair of salivary glands was significantly (p<0.005) higher for *Culex pipiens* Paris than the other temperate population of *Aedes albopictus* Nice. The amount of viral RNA copies was also significantly (p<0.005) higher in *Aedes albopictus* Saint-Benoît than in the *Culex quinquefasciatus* population.

**Fig 5:**
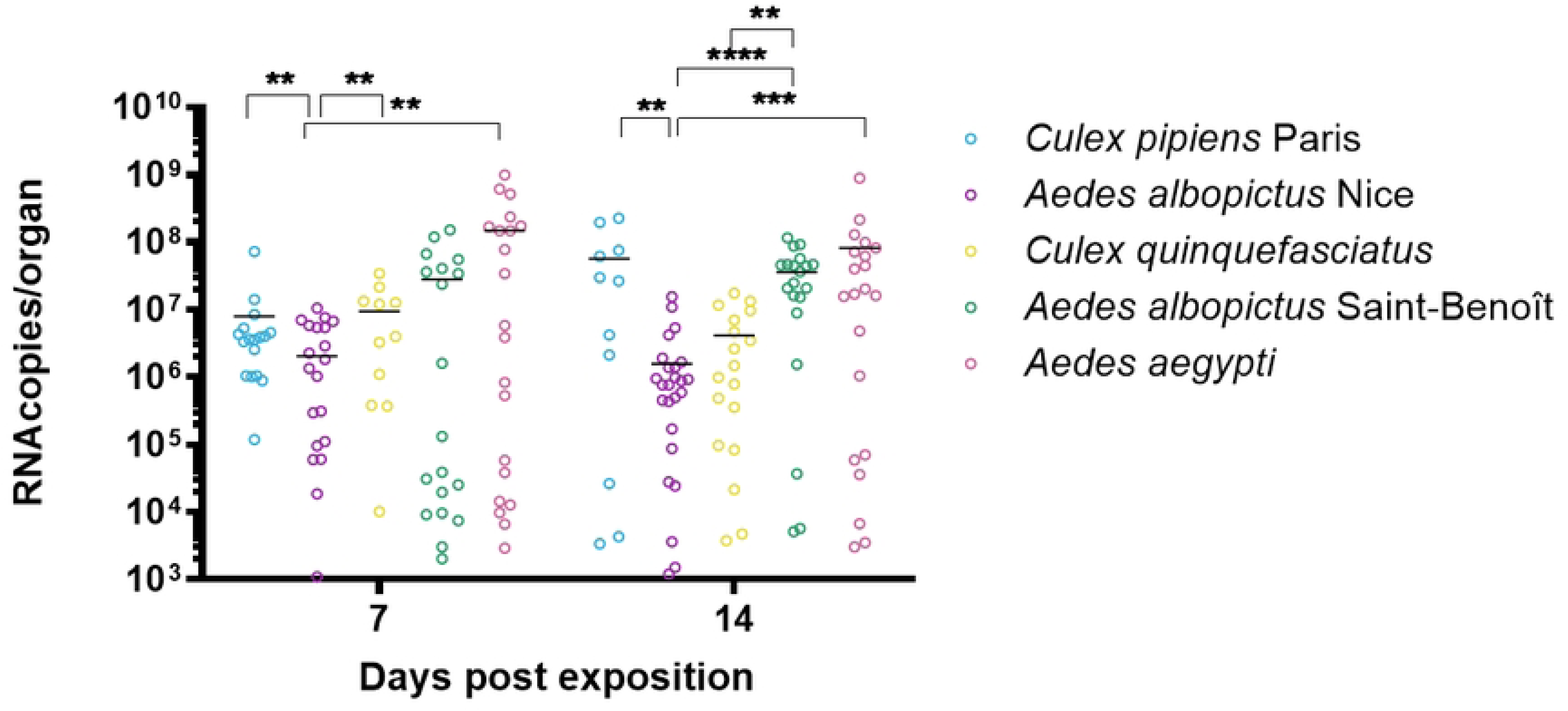
Viral RNA amplification for genome quantification by RT-qPCR in the salivary glands at 7- and 14-dpve. At 7-dpve, n=30 for *Aedes albopictus* Nice and *Aedes aegypti;* n=20 for *Culex quinquefasciatus, Aedes albopictus* Saint-Benoît and *Culex pipiens* Paris. At 14-dpve, n= 20 for *Culex pipiens* Paris and *Aedes albopictus* Saint-Benoit; n=22 for *Aedes aegypti;* n=34 for *Aedes albopictus* Nice and n=40 for *Culex quinquefasciatus.* ns or not represented: non significative; *: p<0.05; **: p<0.01; ***: p<0.001; ****: p<0.0001 (Mann-Whitney non-parametric test)

We could still notice this dichotomy that was more or less important within each species. BBKV viral RNA genomes were detectable at both 7- and 14-dpve in salivary glands of mosquito species.

Viral protein accumulation was also observed in the salivary glands of mosquitoes at 7- and 14-dpve (Fig.1.B). Finding viral proteins there confirms that the virus was replicating in the salivary glands.

These results consolidate that the virus crossed the salivary glands barrier, replicates and produces viral protein.

#### Replication in the salivary glands

Salivary glands were collected (from the same mosquito that were previously used for viral titer analysis) at 7-and 14-dpve and the viral titer was analyzed (Fig 6). No statistical difference was observed at 7-dpve between the different species.

**Fig 6:**
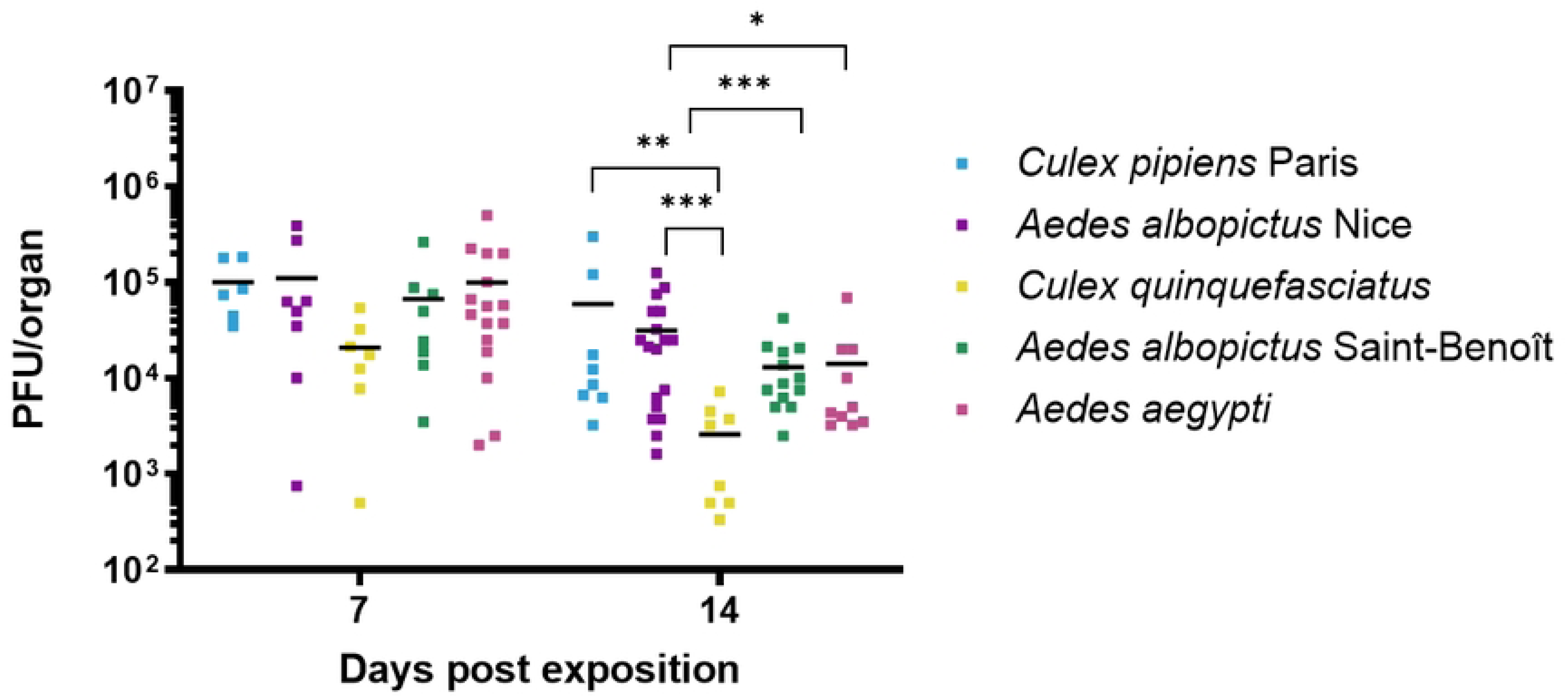
Viral replicative particles quantification in salivary glands. Salivary glands were collected at 7- and 14-dpve. Viral replicative particles were quantified by titration plaque assay on VeroE6 cells. ns or not represented: non significative; *: p<0.05; **: p<0.01; ***:p<0.001; ****: p<0.0001 (Mann-Whitney non-parametric test)

Infectious particles measured in *Culex pipiens*, *Aedes albopictus* Nice, *Aedes albopictus* Saint-Benoît and *Aedes aegypti* salivary glands were similar and statistically higher than those of *Culex quinquefasciatus* at 14-dpve.

A statistical difference in the salivary glands titers was observed between 7- and 14-dpve for *Culex quinquefasciatus, Aedes albopictus* Saint-Benoît *and Aedes aegypti* (p<0.04; p<0.01; p<0.02; p<0.01) for which the viral titers decreased at 14-dpve compared to 7-dpve.

Using this dataset, the dissemination rate to the salivary glands can be determined by the ratio of infected salivary glands to infected midguts (Table 4). At 7-dpve, a statistical difference was observed between all species (p<0.0003), driven by the low dissemination rate in *Aedes albopictus Nice*. No statistical differences were found among the other species. At 14-dpve, no significant differences were observed. However, for *Aedes albopictus Nice*, the dissemination rate was higher at 14-dpve than at 7-dpve (p<0.008), consistent with the increased viral replication in the midgut at this time point.

**Table 4:** Dissemination rate to the salivary glands.

| Mosquito species | Dissemination rate %<br>7-dpve | Dissemination rate %<br>14-dpve |
| --- | --- | --- |
| *Culex pipiens* Paris | 86% (6/7) | 67% (8/12) |
| *Aedes albopictus* Nice | 21% (6/28) | 55% (18/33) |
| *Culex quinquefasciatus* | 70% (7/10) | 57% (8/14) |
| *Aedes albopictus* Saint-Benoît | 59% (10/17) | 76% (13/17) |
| *Aedes aegypti* (PAEA) | 77% (17/22) | 83% (10/12) |

The dissemination rate represents the number of infected salivary glands over the number of positive- infected midguts at 7- and 14-dpve.

#### BBKV transmission through mosquito saliva

Before detecting any viral protein in the saliva, the salivation needs to be confirmed. A first Dot blot on saliva samples was performed using polyclonal antibodies against the saliva protein of *Aedes* or *Culex* species as a primary antibody (Fig 7).

**Fig 7:**
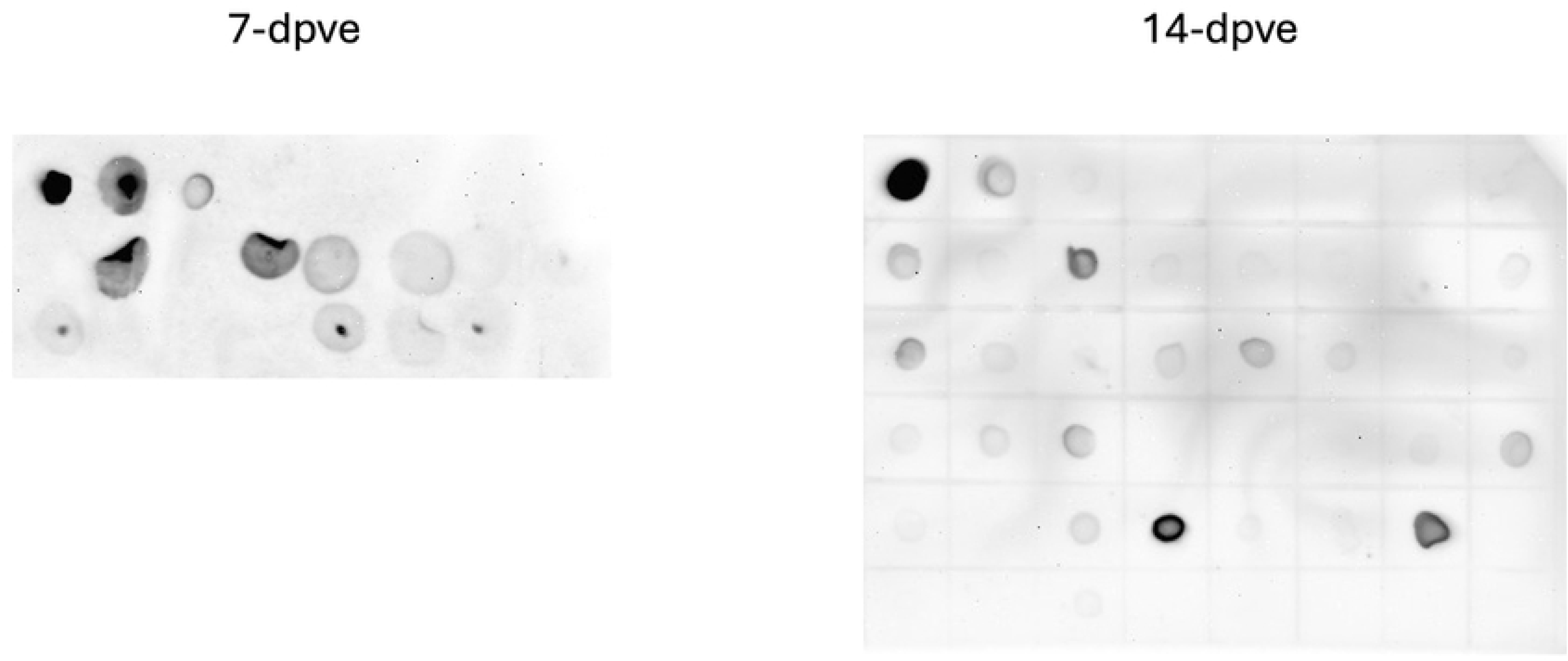
Detection of effective salivation on saliva samples collected. Saliva samples were deposited on PVDF membranes. After drying and blocking with a skimmed milk solution, the membranes were incubated in the presence of an anti-saliva polyclonal antibody and revealed by a peroxidase-labeled secondary antibody. The first row represents the positive controls with fold-dilution and negative controls and the rows below show results obtained with the samples.

We therefore confirmed that most of the mosquito tested had salivated.

We then performed RT-qPCR on saliva samples (Fig 8). We found out that there are viral RNA at both 7- and 14-dpve. The level of RNA copies did not differ between the various species tested.

**Fig 8:**
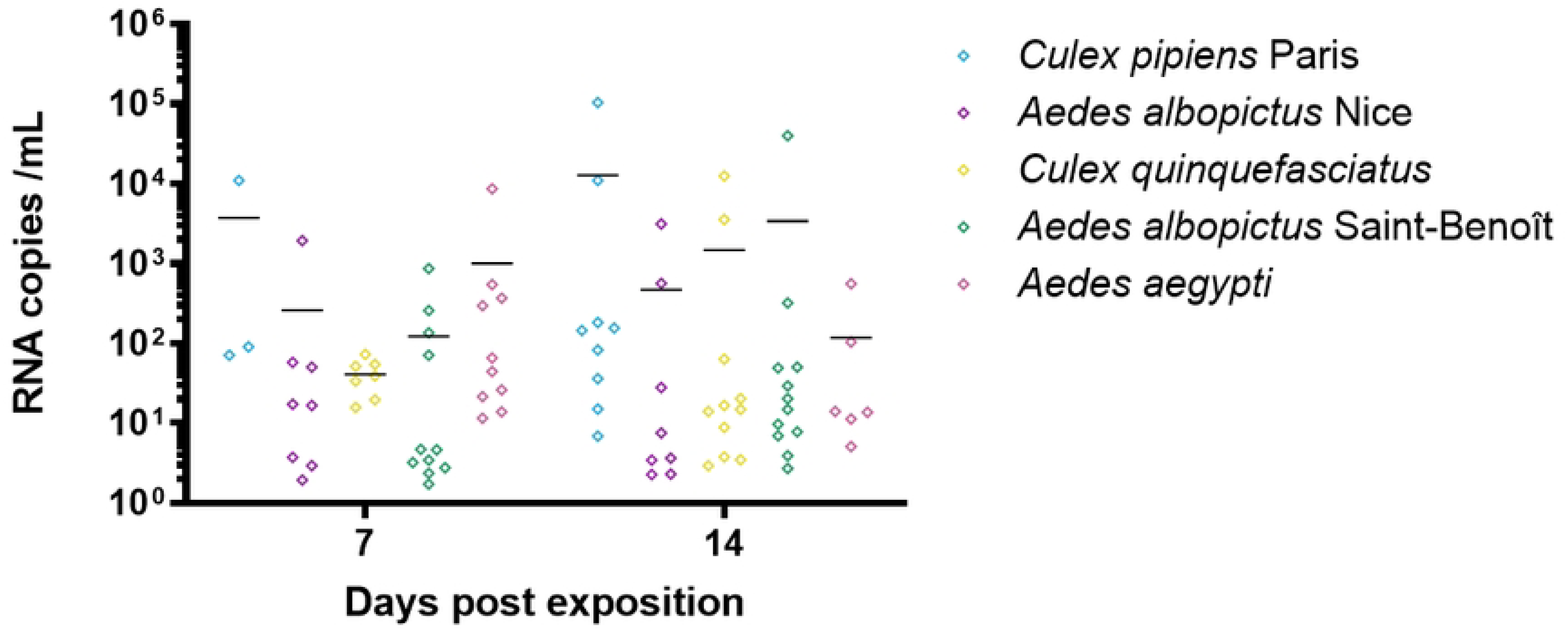
Viral RNA amplified for genome quantification by RT-qPCR in the saliva at 7- and 14-dpve. Saliva samples were collected at 7- and 14-dpve to BBKV. They were submitted to RNA extraction and RT-qPCR. Statistical tests were found not significative between the difference species Kruskal-Wallis non-parametric test)

Titration assays were performed to confirm if infectious viral particles were released in saliva. Viral titers ranging to 10^2^ to 10^3^ pfu were found in the saliva of all species tested (data not shown).

## Discussion

The study provides the first detailed assessment of BBKV vector competence across multiple mosquito species from both temperate and tropical regions. Although isolated from *Mansonia africana* mosquitoes as far back as 1969, BBKV remains poorly characterized, raising concerns due to its association with febrile illness, rash, and arthritis in both humans and animals, with limited treatment options. The findings from this study have significant implications for understanding the potential emergence of BBKV in various regions worldwide through competent mosquito vectors.

To overcome the study and limit variations, a molecular clone of BBKV was designed. Previous studies using molecular clones for Ross River and Chikungunya alphaviruses have demonstrated significant biological research interest (24,25). This molecular clone is a valuable tool and may serve as a foundation for developing other molecular tools, such as replicons, pseudo-particles, and other virus-like particles to address other tropism analysis. Viral stocks were produced from the molecular clone in both insect and mammalian cells, with the insect cell-produced stock chosen based on previous observations regarding SINV (26).

To assess the potential risk of viral BBKV emergence in both tropical and temperate regions, several mosquito species were infected. *A. aegypti* PAEA strain from French Polynesia served as a laboratory control, while *A. albopictus* from Nice (South of France) and *C. pipiens* from Paris (France) represented temperate mosquitoes. *A. albopictus* from St-Benoît (La Réunion) and *C. quinquefasciatus* represented tropical populations. These populations were selected because *Aedes* mosquitoes are anthropophilic while *Culex* species show ambivalence, being both anthropophilic and ornithophilic. Previous studies have highlighted the role of migratory birds in the global dissemination of SINV (27–29).

While *Aedes* mosquitoes are well known for transmitting alphaviruses, *Culex* species are less commonly associated with these viruses. However, SINV can be detected in various vector populations of both *Aedes* and *Culex* species in northern Europe (29,30), Germany (31) and in *C. modestus* in Czech Republic (32,33). In Brazil, *C. quinquefasciatus* was found to be infected by Mayaro virus (MAYV) but was not competent to transmit it (34,35). *C. neavei* has also been described as competent for SINV in La Réunion (36).

Key findings demonstrate vector competence in several mosquito species for BBKV, from both temperate and tropical regions. Infection rates for *Aedes* species consistently exceeded 70%, while *Culex* species exhibiting rates above 40% at 7-dpve. *Aedes* mosquitoes have therefore a higher infection rate than the *Culex* mosquitoes and BBKV replicates better in the midguts of *Aedes* species than *Culex*. Interestingly, the tropical population of *A. albopictus* maintains and amplifies BBKV overtime in the midguts. Regardless of infection rates, viral titers in the midgut were between 10^3^ and 10^5^ PFU/organ for all species at both 7- and 14-dpve. Examination of salivary glands revealed the presence of viral RNA and proteins in both *Culex* and *Aedes* species, confirming BBKV’s ability to cross the salivary gland barrier and replicate within these glands. Infectious and replicative viral particles were found in mosquito saliva, suggesting potential transmission to a new vertebrate host. A recent study demonstrated that after artificial blood meal infected with wild-type BBKV, viral RNA in the bodies of female tropical mosquito species (*Aedes aegypti*, *Culex neavei* and *Culex quinquefasciatus)* reached up to 6 log_10_ at 15-dpve (37). In good agreement, in our study, viral proteins and replicative BBKV virus were detected in the midgut and salivary glands of *Culex quinquefasciatus* and *Aedes aegypti* mosquitoes. The viral genome was found in the midgut, legs, salivary glands, and saliva of mosquitoes. Two distinct pools of viral RNA copies were identified: one with lower titers and the other with higher titers. The lower pool may represent immature or defective viral particles (38), while the higher pool may indicate infectious particles. Viral structural proteins were detected in the midgut and salivary glands, suggesting the presence of mature (infectious) particles. Infectious particles were also detected in the midgut and salivary glands. pE1-E2 envelope glycoproteins showed different molecular weights in midguts and salivary glands of BBKV-infected mosquitoes, suggesting tissue-specific post-translational modifications. This observation aligns with studies on SINV glycoproteins, which exhibit different glycan compositions based on the host and tissue type (39–42).

While BBKV and SINV are transmitted by *Culex* mosquitoes, SINV vector competence is better studied with known vector species and a well-documented transmission cycle than BBKV. In 2022, temperate mosquitoes from Germany were also studied for their vector competence for SINV (43). Although these data cannot be strictly compared due to the differences in the viruses used and the mosquito species collected and studied in different regions, they highlight similar trends.

Generally, *Culex quinquefasciatus* can be infected by both SINV and BBKV, with the virus disseminating to various organs and reaching the salivary glands. Viral infections persist up to 14-dpve. It appears that the viral titer is higher when mosquitoes are infected with BBKV compared to SINV. Moreover, infection and persistence seem more pronounced in *Culex pipiens* infected with BBKV than with SINV. We confirmed that *Aedes aegypti* can be both infected by both wild-type BBKV and its molecular clone while *Aedes albopictus* from Europe can be infected by both SINV and BBKV. Viral infections persist over time in these species.

Overall, our findings indicate that mosquito species from both tropical and temperate regions are susceptible to BBKV infection, and that *Culex* mosquitoes are more susceptible to BBKV than to SINV, showing therefore that the two viruses have a different behavior in certain mosquito species.

The ability of BBKV to infect both anthropophilic (*Aedes*) and ornithophilic (*Culex*) mosquitoes raises concerns about its potential for both urban and sylvatic transmission cycles. This broad vector competence suggests that BBKV could be maintained in diverse ecological settings, facilitating its spread across temperate and tropical regions. The virus ability to persist in mosquito populations and replicate efficiently increases the risk of human outbreaks, making it a significant public health concern.

The emergence of alphaviruses is often driven by climate change, urbanization, and deforestation, which alter mosquito distribution and viral transmission dynamics. Given that no specific antiviral treatments or vaccines exist for BBKV, proactive measures are essential. Enhanced arbovirus surveillance programs should be implemented to monitor BBKV circulation in both mosquito populations and sentinel animals, particularly in regions with competent vectors. Additionally, developing diagnostic tools, serosurveillance programs, and risk assessment models will be critical to predicting and mitigating potential outbreaks.

The variability in viral replication among different mosquito species underscores the importance of understanding local vector populations and their role in transmission dynamics. Identifying species-specific differences in infection, dissemination, and transmission efficiency will help refine vector control strategies and public health preparedness efforts.

Future research should focus on elucidating the precise transmission mechanisms and developing effective strategies for surveillance, prevention, and control.

## Acknowledgments

The authors are grateful to Carine Maisse-Paridisi for her help in designing the Babanki virus primer. The authors are also grateful to Mylène Weill for kindly providing *Culex quinquefasciatus* (Slab strain) larvae.

## Authors contributions

Mathilde Ban: Conceptualization, Formal Analysis, Investigation, Methodology, Vizualisation, Writing – Original Draft Preparation, Writing – Review & Editing

Virginie Geolier: Investigation

Elisabeth Ferquel: Formal analysis, Writing – Review

Martin Faye: Methodology, Resources, Software, Writing – Review & Editing

Moussa Moise Diagne: Methodology, Resources, Software, Writing – Review & Editing

El Hadji Ndiaye: Methodology

Gamou Fall: Resources, Project administration, Writing – Review & Editing

Mawlouth Diallo: Methodology, Resources

Dimitri Lavillette: Conceptualization, Resources, Supervision, Writing – Review & Editing

Valérie Choumet: Conceptualization, Resources, Supervision, Validation, Vizualisation,

Writing – Original Draft Preparation, Writing – Review & Editing

